# The endometrial transcription landscape of MRKH syndrome

**DOI:** 10.1101/2020.02.18.954768

**Authors:** T Hentrich, A Koch, N Weber, A Kilzheimer, S Burkhardt, K Rall, N Casadei, O Kohlbacher, O Riess, JM Schulze-Hentrich, SY Brucker

## Abstract

The Mayer-Rokitansky-Küster-Hauser (MRKH) syndrome (OMIM 277000) is characterized by agenesis of the uterus and upper part of the vagina in females with normal ovarian function. While genetic causes have been identified for a small subset of patients and epigenetic mechanisms presumably contribute to the pathogenic unfolding, too, the etiology of the syndrome has remained largely enigmatic. A comprehensive understanding of gene activity in the context of the disease is crucial to identify etiological components and their potential interplay. So far, this understanding is lacking, primarily due to the scarcity of samples and suitable tissue.

In order to close this gap, we profiled endometrial tissue of uterus rudiments in a large cohort of MRKH patients using RNA-seq and thereby provide a genome-wide view on the altered transcription landscape of the MRKH syndrome. Differential and co-expression analyses of the data identified cellular processes and candidate genes that converge on a core network of interconnected regulators that emerge as pivotal for the perturbed expression space. With these results and browsable access to the rich data through an online tool we seek to accelerate research to unravel the underlying biology of this syndrome.

## Introduction

Mayer-Rokitansky-Küster-Hauser (MRKH) syndrome [OMIM 277000] is the second most common cause of primary amenorrhea with an incidence rate of about one in 4000 to 5000 female births (1). It is defined by agenesis of the uterus and the upper part of the vagina in 46, XX females with normal ovarian function and normal secondary sexual characteristics. The syndrome may occur either in an isolated form (type 1) or in association with extragenital abnormalities (type 2) such as renal or skeletal malformations (2, 3).

The spectrum of malformation encountered in MRKH patients suggests the disease to originate from a developmental defect of the intermediate mesoderm during embryogenesis, yet the etiology of the syndrome remains largely enigmatic. While most cases are sporadic, familial cases exist and imply a genetic component in the etiology (4-6). Specifically, chromosomal aberrations in 1q21.1, 16p11.2, 17q12, and 22q11 as well as mutations in *LHX1, TBX6, RBM8A*, and *WNT9B* have been linked to MRKH. Additionally, mutations of *WNT4* cause an atypical form of the syndrome characterized by hyperandrogenism (7).

*LHX1, WNT4*, and *WNT9B* play important roles in the formation of the Müllerian Ducts (MD) from the coelomic epithelium in gestational week six (8, 9). The freshly formed MDs start growing caudally along the Wolffian Ducts. By week eight, both MDs begin to fuse and make contact with the uterovaginal sinus. In males, the MDs start to regress after week ten under the influence of *AMH* and *WNT7A*. In females, however, they differentiate into ovaries, uterus, cervix, and vagina under control of *ESR1, HOXA* and *WNT* genes. In this context, *HOXA9, HOXA10, HOXA11*, and *HOXA13* are essential for correct tissue patterning. Their expression is tightly controlled through *Wnt signalling* and histone methylation marks (10-12) suggesting epigenetic principles to also play a role in the unfolding of the disease.

Towards a better understanding of the etiology, examining perturbed gene activity on a genome-wide scale promises to identify regulatory hubs on which genetic or epigenetic contributions converge. Attempts to identify the molecular mechanisms of the syndrome have been hampered by the lack of a comprehensive transcriptome profile for primary tissue in MRKH patients. This obstacle can partly be attributed to the fact that patients do not always have uterus rudiments with a complete endometrial layer and to the scarcity of uterine tissue resulting from challenging collection and biobanking efforts.

In order to close this gap, we have assembled a large and unique cohort of MRKH type 1 and type 2 patients and profiled the transcriptome in endometrial tissue. The expression landscape that emerged along comprehensive differential and co-expression analyses of these data mapped known and novel candidate genes and identified regulatory networks that seemingly drive the underlying disease biology. By offering an online tool that allows navigating and downloading these rich data from single genes to pathways, we seek to provide a much-needed building block for the research community to understand the molecular pathomechanisms of MRKH.

## Results

### Widespread transcriptome changes in endometrial tissue of MRKH patients

To investigate disease-associated perturbations to the endometrial transcriptome of MRKH patients, we performed RNA-seq of uterine rudiments obtained from 39 patients (22 type 1 and 17 type 2) as well as 30 controls. Throughout the analysis pipeline, stringent quality filters were applied, which also helped to identify a few outlier samples using clustering techniques (Supplementary Fig. 1A, B).

As no obvious sequencing parameters differed for these samples and no batch effects were apparent, tissue composition differences resulting from the sample collection process were considered. After integration of the data with single-cell data from endometrial tissue that recently became available (13), the expression signature of cell type-specific markers, in particular for ciliated and unciliated epithelial cells, indeed distinguish the outlier samples from all other samples (Supplementary Fig. 1C) and points to a different underlying cell type composition. Since it is difficult to assess the combinatorial complexity and effects of these convolutions computationally, the affected samples were removed from all subsequent analyses, which left a total of 60 high-quality samples with consistent expression signatures (Supplementary Fig. 2).

In a first step, differential expression changes were determined between MRKH patients and control samples. According to thresholds of *p*_BH_ ≤ 0.05 and |log_2_*FC*|≥ 0.5, a total of 1906 differentially expressed genes (DEGs) comprising 1236 up- and 670 downregulated genes in MRKH type 1 and 1174 DEGs with 801 up- and 373 downregulated genes in MRKH type 2 were identified when compared to controls (Fig. 1A). These numbers of affected genes in each disease type indicate profound transcriptome changes in the endometrium of MRKH patients.

**Figure 1:**
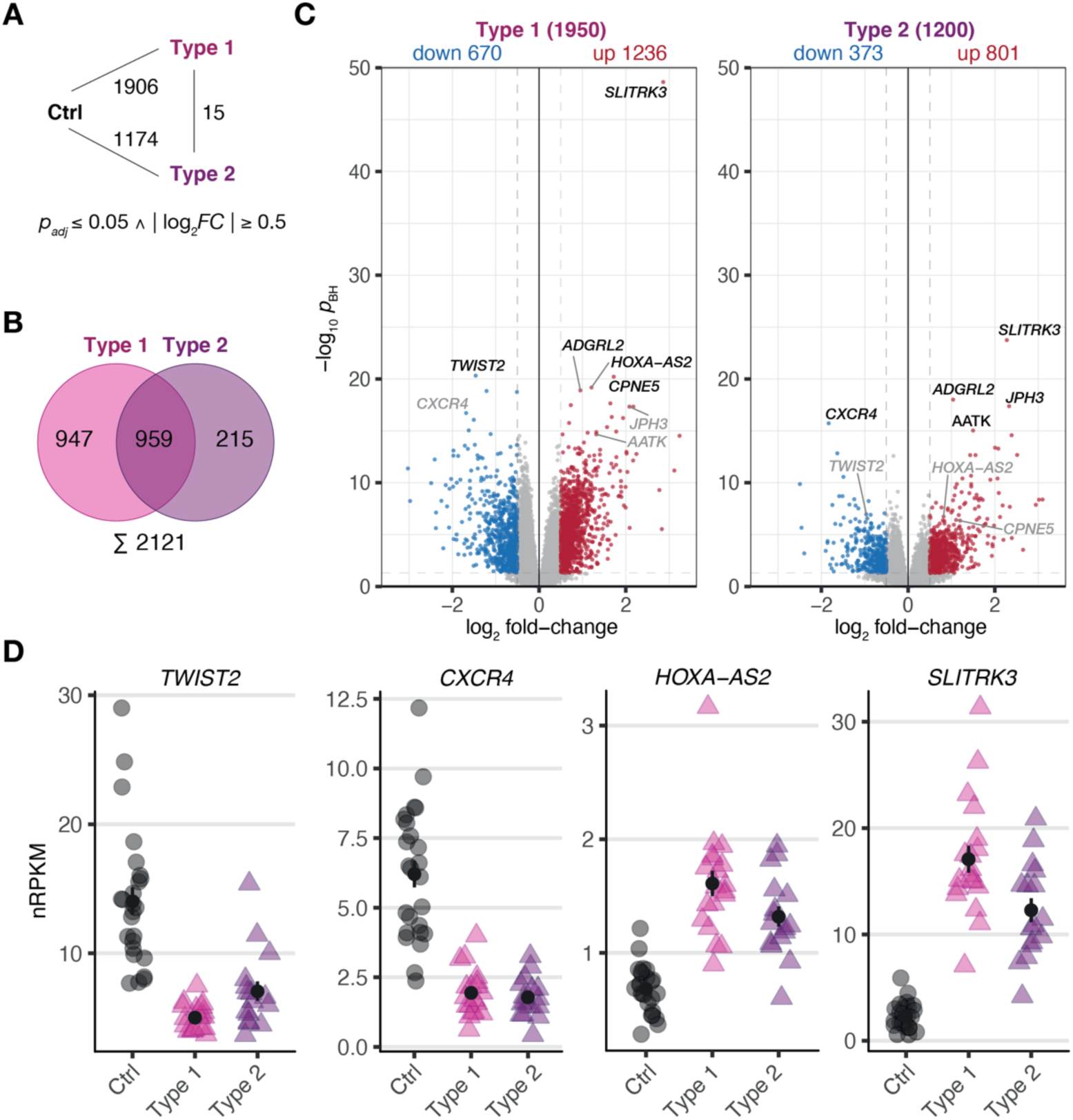
MRKH patients exhibit wide-spread gene expression changes in endometrial tissue compared to unaffected controls. **(A)** Schematic diagram of three experimental groups (Ctrl, unaffected women; Type 1, patients with MRKH type 1; and Type 2, patients with MRKH type 2) indicating number of differentially expressed genes (DEGs) for each pairwise comparison. Fold change and significance cut-offs below. **(B)** Venn diagram comparing common and distinct DEGs between MRKH type 1 and 2. **(C)** Volcano plot showing magnitude and significance values of gene expression changes identified in endometrial tissue of MRKH type 1 (left panel) and MRKH type 2 patients (right panel) compared to unaffected controls. Blue and red dots highlight DEGs according to the applied cut-offs (see a). The five most significant DEGs from each comparison are labeled. Grey gene labels indicate location of DEGs from the opposite comparison. **(D)** Expression levels for the most significant up- and downregulated DEGs as well as the lincRNA *HOXA-A2* plotted as individual data points with mean ± SEM.

### Largely similar endometrial expression profiles in MRKH type 1 and 2 patients

Next, the DEG sets of each pairwise contrast were compared in order to better understand common and distinct expression changes for the disease subtypes. While overlapping the DEGs by name, about half of them first seemed exclusive for type 1 or type 2, respectively (Fig. 1B). Directly contrasting the subtypes in the differential analysis, however, identified only 15 DEGs (Fig. 1A, Supplementary Fig. 3), which suggested largely comparable perturbations in gene activity in type 1 and type 2.

Despite similar magnitudes of expression changes in both disease types, affected genes in type 2 samples separated less significantly from controls (Fig. 1C), pointing to a larger variability among type 2 samples and coinciding with the greater heterogeneity of clinical features in this disease type. Yet, the localization of the top most significant DEGs in the volcano plot (Fig. 1C) as well as the correlation of expression changes (spearman rank, *r* = 0.87) point at stark similarities between both types. Indeed, the most significant DEGs showed nearly identical expression changes on a gene and transcript isoform level in both disease types (Fig. 1D, Supplementary Fig. 4). This high degree of concordance also becomes apparent from the per-sample expression profiles for the union of all DEGs (Fig. 2). Further, the expression changes were comparable between sporadic cases and patients from families with more than one affected sibling (Fig. 2). Hence, all subsequent analyses were based on the genes underlying this perturbance signature.

**Figure 2:**
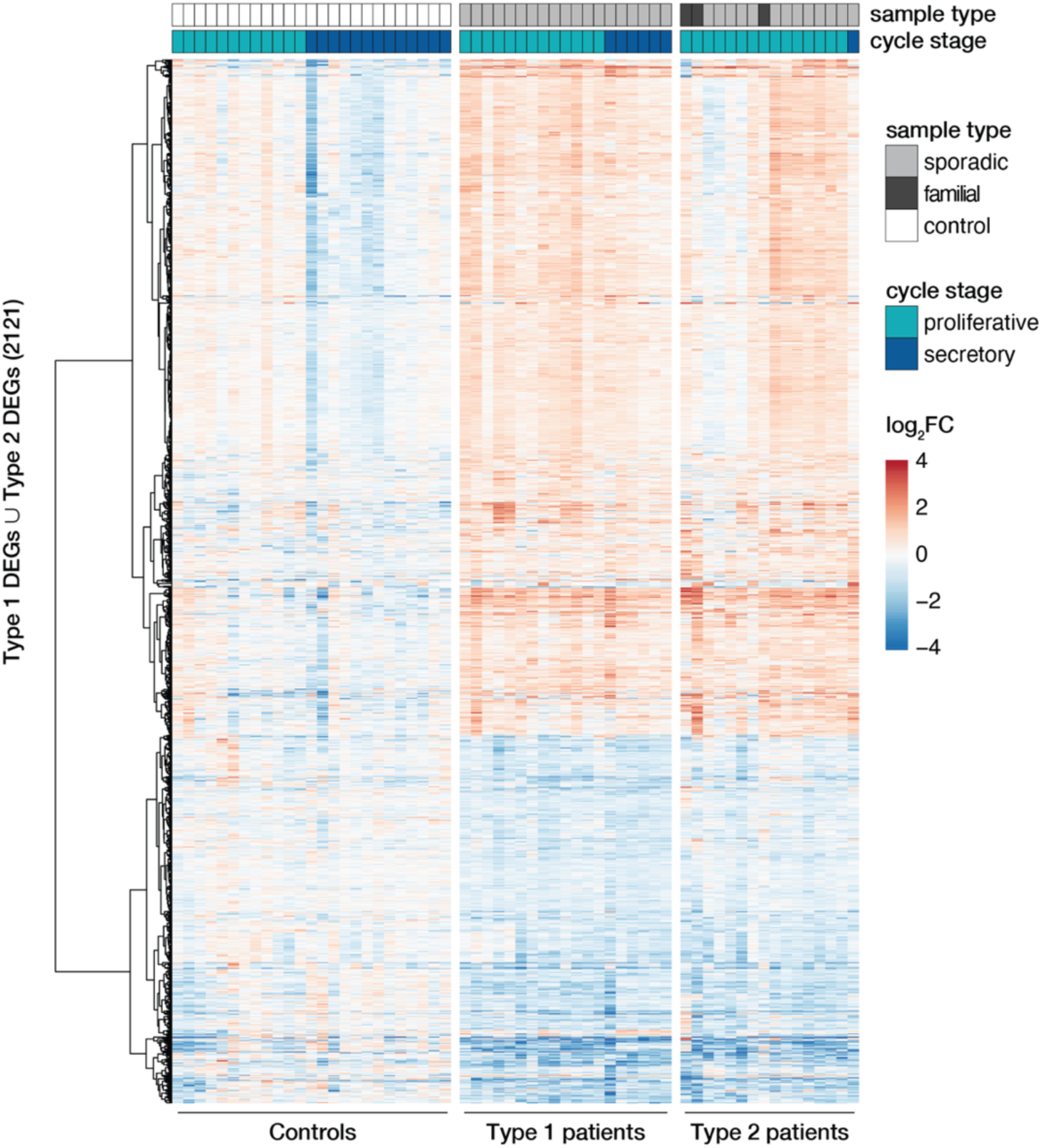
MRKH type 1 and 2 patients show largely similar perturbation patterns in endometrial gene expression. Expression profiles (*log*_*2*_ expression change relative to Ctrl group) of 2121 DEGs (union of DEGs indicated in Fig. 1B) across all samples. Rows hierarchically clustered by Euclidian distance and *ward.D2* method. Cycle information (proliferative or secretory) and patient type (sporadic, familial, or control) on top. For details see Supplementary Table 1.

### Endometrial gene expression changes during the menstrual cycle are disrupted in MRKH patients

Upon closer inspection of the perturbation signature, the heatmap also shows patterning between the proliferative and secretory cycle stage in control samples for a subgroup of genes (upper part of Fig. 2). In MRKH patients, however, this menstrual cycle dependency seems to be largely lost. To better quantify this observation, we determined differential expression between the proliferative and secretory phase in control samples, which yielded 818 DEGs (Supplementary Fig. 5A). Their associated gene ontology (GO) terms were enriched most significantly for *collagen-containing extracellular matrix* (Supplementary Fig. 5B), agreeing with remodeling processes of the extracellular matrix along the transitions between cycle stages (14). In contrast, only 116 genes were identified as cycle-dependent in MRKH type 1 (Supplementary Fig. 5A), indicating that cyclic expression adaptations were damped or lost entirely in these patients despite normal hormone profiles (Supplementary Table 1). Instead, the expression of cycle-dependent genes seemingly remained in the proliferative phase throughout the menstrual cycle (Supplementary Fig. 5C). The analogous analysis for type 2 was omitted due to the highly skewed sample distribution with respect to cycle stages. Together, these analyses are in line with previous reports that the endometrium of MRKH patients does not respond correctly to cycle hormones (15-18).

### Transcriptome changes point to regulators of cell adhesion and development

To unravel the underlying biology of the endometrial MRKH signature, enrichment analyses were applied to identify potential key regulators as well as affected pathways and cellular processes. With respect to GO terms, *plasma membrane part* was the most overrepresented cellular comportment, and *cell adhesion* and *biological adhesion* emerged as most significant biological processes followed by *anatomical structure development* (Fig. 3A).

**Figure 3:**
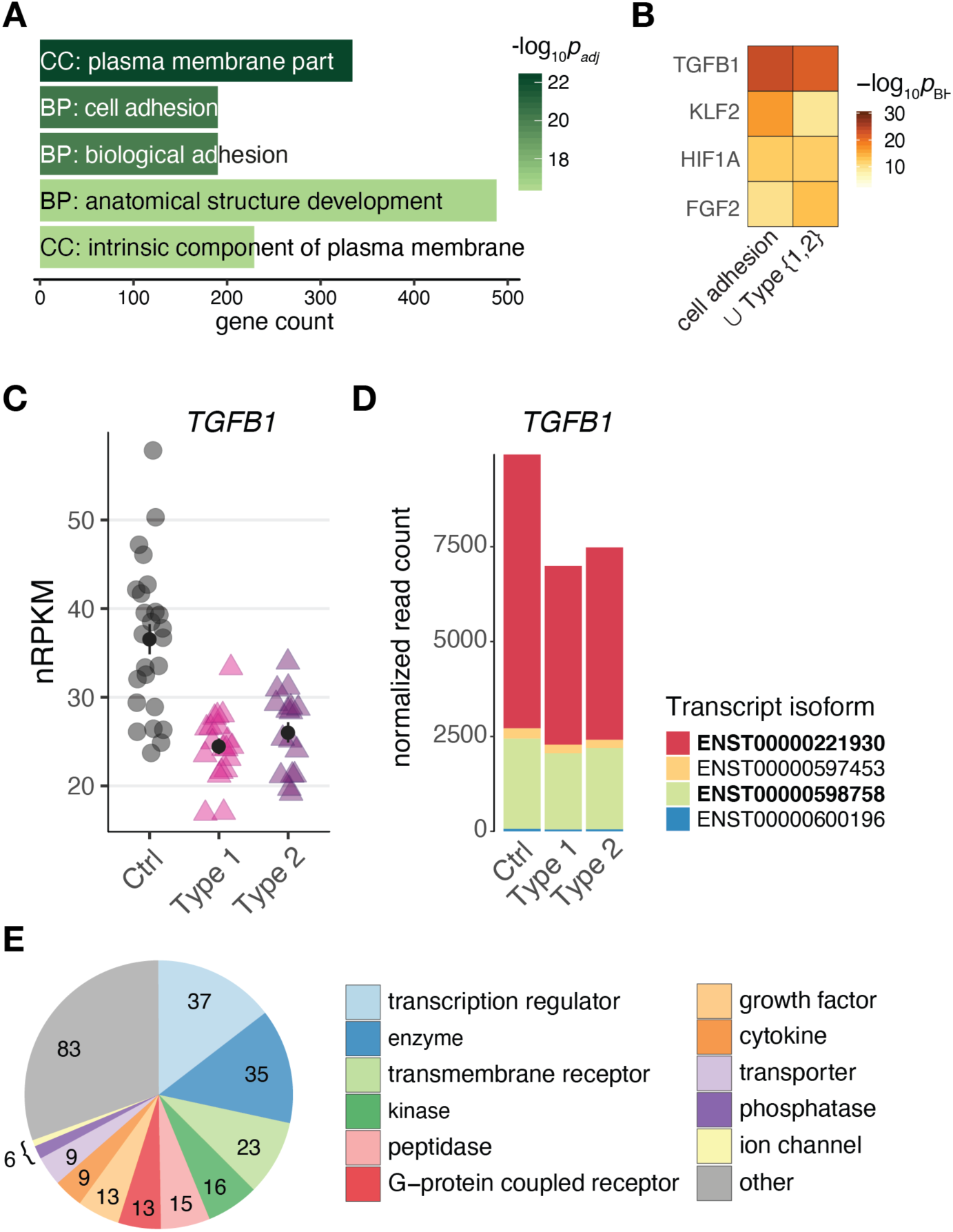
Changes of gene expression in both types of MRKH point to regulators of cell adhesion and development. **(A)** Enrichment analysis identified several significantly overrepresented Gene Ontology terms among the 2121 DEGs (union indicated in Fig. 1B). Top five terms with number of associated genes are shown according to their significance. CC: cellular compartment, BP: biological process. **(B)** Comparison of predicted upstream regulators for the DEGs underlying the cell adhesion term (see A) as well as all 2121 DEGs (union from Fig. 1B) based on Ingenuity Pathway Analysis. Top three significant regulators for each gene set shown. **(C)** Expression changes for TGFB1 plotted as individual data points with mean ± SEM. **(D)** Transcript isoform-specific expression changes of TGFB1 across all conditions. Mean normalized read counts plotted; bold isoforms are protein-coding. **(E)** Among the 253 predicted interactors of TGFB1 differentially expressed in both types of MRKH, transcriptional regulators represent a largest subgroup. Interactors identified based on Ingenuity Pathway Analysis.

Based on binding-site analyses, motifs of the differentially expressed transcription factors *EGR1* and *KLF9* were most significantly overrepresented among the DEGs (Supplementary Fig. 6 A, B). In addition, approaches that integrate ChIP-seq data into such analyses and thereby account also for indirect binding events and factors with less clear motifs (19), suggested the DEGs to be highly enriched for *EZH2* targets (Supplementary Fig. 6C, D). EZH2 (Enhancer of zeste homolog 2), a histone methyltransferase and a catalytic component of PRC2, showed a trend towards up-regulation in MRKH patients (Supplementary Fig. 6E).

To extend the transcription factor-centered analyses to other regulatory mechanisms underlying the observed gene expression changes, we used curated interactome data and mined for regulatory enrichments. From these analyses, *TGFB1* was predicted to be the top upstream regulator for the entire DEG set as well as for the subset of DEGs underlying *cell adhesion* as the most likely affected biological process (Fig. 3B). *TGFB1* showed a down-regulation that resulted predominantly from the longer protein-coding transcript isoform (Fig. 3 C, D). Intriguingly, TGFB1 is known to interact not only with EGR1, KLF9, and EZH2, but also connects to more than ten percent of all DEGs (253 of 2121), many with regulatory capacity, too (Fig. 3E). These results hint at the regulatory neighborhood of TGFB1 as a key modulator of gene expression changes in MRKH.

### Co-expression analysis ranked disease relevance of TGFB1 interactors

To further assess the regulatory relevance of the TGFB1 neighborhood identified along the differential expression analysis, in the next step, a co-expression approach was employed in order to capture groups of genes that change and often function together (20).

Partitioning of the perturbed endometrial expression space using weighted correlation network analysis (21) led to 35 co-expression modules that ranged from 39 to 3,268 genes in size and totaled to 15,361 genes (Supplementary Fig. 7A). In this manner, the co-expression analysis reduced thousands of genes to a relatively small number of coherent modules that represent distinct transcriptional responses. To quantify the overall relationship between modules and the disease, correlations with module eigengenes (summary expression profiles) were calculated (21). After filtering and correcting with *p*_BH_ ≤ 0.05 and Bayes factor ≥ 3, twenty modules (six up- and 14 downregulated) passed the significance cut-off (Fig. 4A and Supplementary Fig. 7B). Furthermore, the meta-analysis significance statistics ranked the modules by their overall association with the disease (Fig. 4A) and yielded a measure of module membership for all genes in all modules. The module membership measures how similar the gene expression profile is to a module’s eigengene. Genes whose profiles are highly similar to the eigengene are considered hub genes and have been shown to implicate relevant biological functions (21).

**Figure 4:**
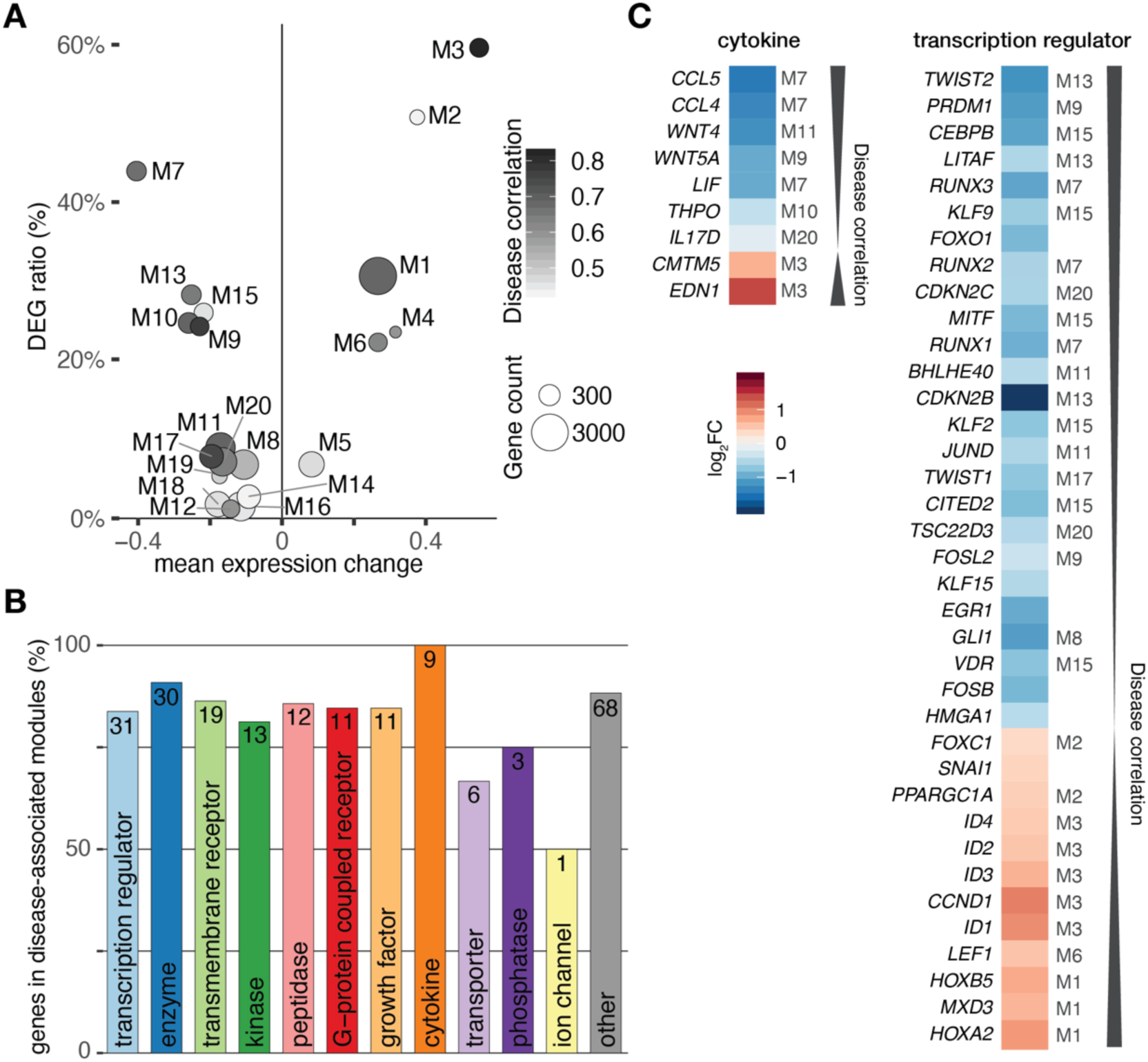
Interactors of TGFB1 reach in all disease-associated co-expression modules. **(A)** Weighted gene correlation network analysis (WGCNA) identified 35 co-expression modules of which 20 were significantly associated with the disease. **(B)** Bar diagram depicting the number of TGFB1 interactors in disease-associated modules. Absolut number within bar as well as amount in percent shown on y-axis for each functional type. **(C)** Cytokines and transcriptional regulators predicted to interact with TGFB1 are in highly disease-associated co-expression modules. Module of interactors indicated for significant modules only.

According to these characteristics, *TGFB1* located to the disease-associated module M13, correlated significantly with the disease (*r* = 0.68, *p* ≈ 10^−9^), and was among the top 50 hub genes of this module. Of the 253 TGFB1 interactors, 214 reached into all 20 significant disease-associated modules (Fig. 4B). Genes annotated for *transcriptional regulator* constituted the largest subgroup of interactors, accompanied by all interacting *cytokines* found in high-ranking modules (Fig. 4B, C). Among them were *WNT4* and *WNT5A* of the WNT signaling pathway as well as *HOXA2* and *HOXB5* as members of the HOX clusters (Fig. 4C, Supplementary Fig. 8), all of which have been associated with MRKH (7). In addition, *TWIST2* identified as one of the genes with the most significant expression change (Fig. 1D) ranked highest among the interacting transcription regulators.

### Transcriptome changes in MRKH converge on regulatory loops in the TGFB1 neighborhood

The combined approach of differential and co-expression analyses highlighted candidate genes that can explain large parts of the altered expression landscape in context of the MRKH syndrome. These novel candidates together with previously associated genes like *FOXO1* (22) and pathways like *WNT signaling* (7) share direct links into the TGFB1 regulatory neighborhood (23) and reach into disease-associated co-expression modules.

Intriguingly, many interactors are not only targets of TGFB1, but often also upstream regulators, hence, forming regulatory loops (Fig. 5). Along such loops, the predicted transcriptional regulators EGR1, KLF9, and EZH2 are found, too. These loops are connected to a dense core network that emerges as pivotal in explaining the disease signature and comprises some of the most significantly altered genes with potent regulator capacity like *TWIST2*.

**Figure 5:**
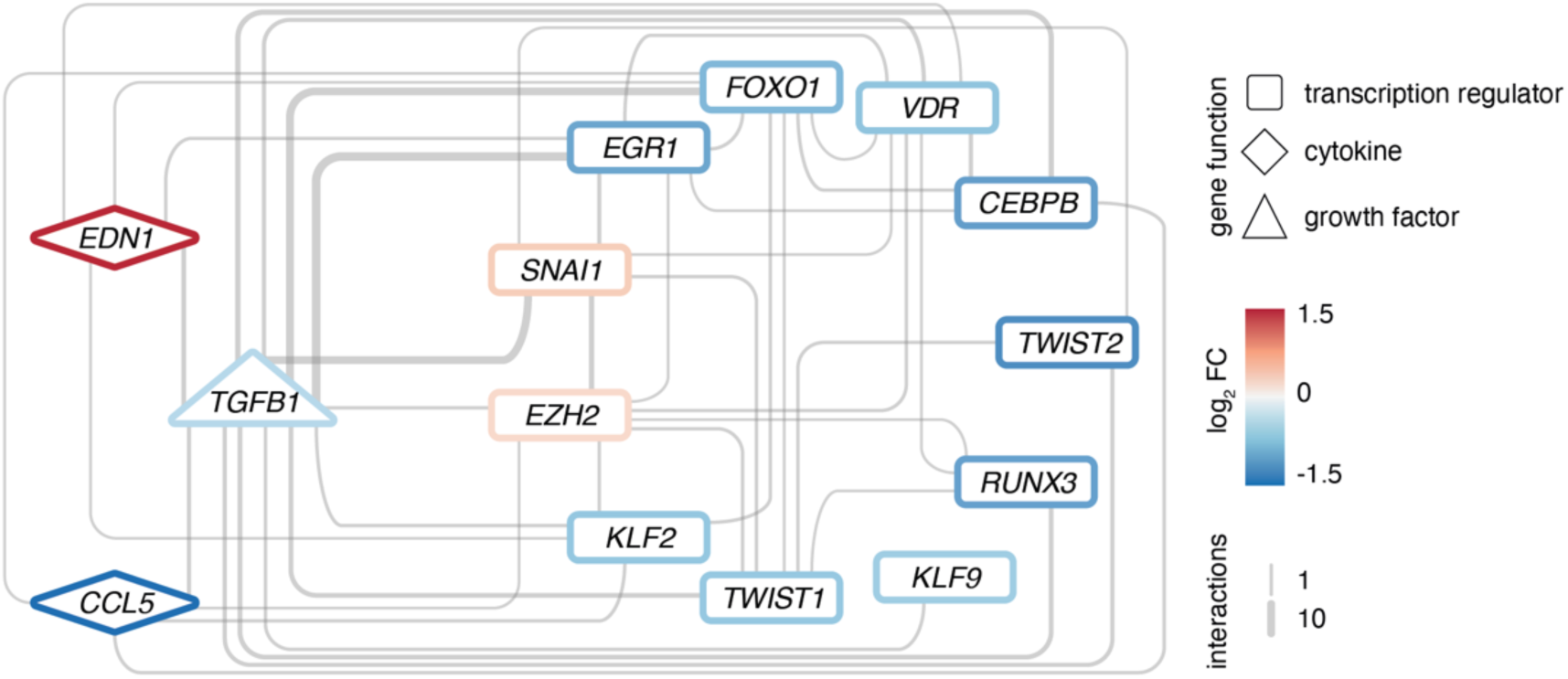
Regulatory loops around TGFB1 link important transcription factors and cytokines. Network of EZH2 as well as upstream interactors of TGFB1 among the 2121 DEGs, respectively. Interactions based on Ingenuity Pathway Analysis and filtered for transcription regulators and cytokines. All interconnections between genes shown. Genes color-coded by mean expression change observed in MRKH / Ctrl. Line width indicates number of curated interactions.

To further disentangle the regulatory relations in the core network towards a potential point of origin, information from other regulatory layers or functional experiments is required. With respect to the former, epigenomic interrogations might yield additional insight given the prominent location of EZH2 and its role in development. With respect to the latter, the network might serve as starting point to select candidate genes for functional characterizations. To facilitate the selection process and put choices into perspective with respect to other gene expression changes, we offer an online tool that allows downloading, visualizing, and navigating through the endometrial transcription landscape from single genes to entire pathways that can be accessed here: http://mrkh-data.informatik.uni-tuebingen.de.

## Discussion

In this study, we assembled a large cohort of patients with type 1 and type 2 MRKH syndrome and profiled the endometrial transcriptome. The key goal of these efforts was to gain a genome-wide understanding of expression changes in order to identify dysregulations and potential origins of the disease, as only a fraction of MRKH cases can be traced to genetic defects.

Our analyses first revealed widespread perturbations of gene activity in the endometrium that were highly similar between type 1 and type 2 patients. The genes underlying this shared perturbance signature point to key regulators that are centrally linked to cell adhesion and developmental processes. The observed expression similarity between type 1 and type 2 cases agrees with previous microarray interrogations of myometrial tissue (24, 25).

Despite highly similar expression perturbations, phenotypically MRKH type 1 and type 2 patients differ. Type 1 cases are characterized by utero-vaginal malformation only, while type 2 patients display a more complex phenotype that entails non-genital abnormalities. Specifically, the urogenital tract including the kidneys is frequently affected in type 2 (e.g. unilateral kidney agenesis, ectopia of one or both kidneys, and horseshoe kidneys). Furthermore, skeletal anomalies, hearing defects, cardiac, and digital anomalies as well as ciliopathies occur in type 2 cases (3). The utero-vaginal malformations, however, are highly similar between both disease types. Uterus rudiments exist in both, although to a lesser extent in type 2.

As the innermost lining layer of the uterus, the endometrium consists of multiple cell types in a basal and functional tissue layer. As the latter thickens and is shed during menstruation, the endometrium undergoes substantial modifications during the proliferative, secretory, and menstrual phase. The correct staging of these phases is governed by cyclic gene activity over the course of the menstrual cycle (14). In line, we observed expression changes between the proliferative and secretory phase in control samples. Intriguingly, these were largely lost in MRKH patients. Instead, the expression of most genes remained in the proliferative phase although the hormonal profiles indicate patients were in the secretory phase. This finding agrees with previous studies that describe lacking responsiveness of the endometrium to hormones in MRKH patients (15-18). The transcriptome data we provide now offer the opportunity to trace the phenomenon to individual genes and pathways and examine co-occurring effects.

Developmentally, the uterus as well as the upper two thirds of the vagina originate from fusion of the Muellerian ducts. In context of the MRKH syndrome, this fusion seems inhibited in gestational week eight and only two uterine rudiments and a vaginal dimple are formed (18). They remain in this incomplete embryonic stage and do not undergo normal enlargement at the beginning of adolescence. As the malformation manifests early during embryonic development, associated pathways have been proposed to be key for MRKH syndrome. In keeping, we identified significant enrichments for *cell adhesion* and *anatomical structure development* among perturbed genes. In addition, developmental regulators like *TGFB1* and *EZH2* emerged as central from the analyses.

*TGFB1* was significantly downregulated in MRKH patients and belongs to the superfamily of transforming growth factor β (TGFβ), which is centrally involved in cell growth and differentiation as well as in regulation of female reproduction and development (26). While the uterus of *Tgfb1* mutant mice are morphologically normal, embryos become arrested in the morula stage (27), suggesting critical roles of this gene. Furthermore, TGFβ signaling is crucial for the epithelial to mesenchymal transition (EMT), in which cells lose their epithelial characteristics and acquire migratory behavior (28). EMT is necessary for the development and normal functioning of female reproductive organs such as the ovaries and the uterus and dysregulation may cause endometriosis, adenomyosis, and carcinogenesis (29).

TGFB1 is linked to Enhancer of zeste homolog 2 (EZH2) (30-32), the most overrepresented transcriptional regulator predicted to bind to the DEGs according to motif analysis and ChIP-seq reference data. EZH2 is the rate-limiting catalytic subunit of the polycomb repressive complex 2 that silences gene activity epigenetically through deposition of the repressive H3K27me3 histone mark (33).

In MRKH patients, *EZH2* showed a small but significant trend of upregulation, potentially remains of elevated activity earlier in life. If true, altered levels of EZH2 might have led to falsely deposited H3K27me3 marks in the genome during development which caused perturbations in gene activity and interfered with correct unfolding of the developmental program. The observed transcriptional perturbances at the time of profiling might hence be direct consequences or indirect adaptation attempts of the system.

In mice, uterine EZH2 expression is developmentally and hormonally regulated, and its loss leads to aberrant uterine epithelial proliferation, uterine hypertrophy, and cystic endometrial hyperplasia (34). Furthermore, reduction of EZH2 and ultimately H3K27me3 levels result in increased expression of estrogen-responsive genes (35).

In this context, exposure to environmental estrogens has also been proposed to reprogram the epigenome by inducing non-genomic ER signaling via the phosphotidylinositol-3-kinase (PI3K) pathway (36). The kinase AKT/PKB phosphorylates and inactivates EZH2 and thereby decreases H3K27me3 levels in the developing uterus. Consequently, estrogen-responsive genes become hypersensitive to estrogen in adulthood and cause hormone-dependent tumors to develop. Our results suggest the opposite effect might play a role in MRKH and failure of enlargement in organ size is a consequence of elevated EZH2 levels.

Taken together and given that only a fraction of MRKH syndrome cases can be explained by genetic defects, these hints towards epigenetic dysregulation playing a potential role in the etiology should be further investigated. Towards these efforts, we consider our results and data to serve as reference point and resource for further exploration.

## Methods

### Patient cohort

Endometrial samples were prospectively collected at the Department of Obstetrics and Gynaecology of the University of Tübingen from rudimentary uterine tissue from patients with MRKH syndrome and uterine tissue from healthy controls. Tissue was taken from 39 patients with MRKH syndrome (22 MRKH type 1 and 17 MRKH type 2, see Supplementary Table 1 and 2) at the time of laparoscopically assisted creation of a neovagina (37). As control group, 30 premenopausal patients, less than 38 years of age, who underwent hysterectomy for benign disease, were included in the study (Supplementary Table 1 and 2). Samples were examined histologically and found to predominantly contain endometrial tissue without excluding myometrial residuals. Correlation with the individual cycle phase was achieved by taking standardized histories and by using hormone profiles from peripheral blood taken 1 day before surgery (see below). The study received prior approval by the Ethics Committee of the Eberhard-Karls-University of Tübingen (Ethical approval AZ 397/2006, Nr.28/2008BO1, 205/2014BO1).

### Hormone levels and correlation with cycle phase

Whole blood was taken from patients and controls one day before or after surgery. Blood serum was used to measure LH, FSH, P, and E2 with a chemiluminescence immunoassay (Vitros eci; Diagnostic Product Cooperation). Cycle phase 1 (proliferative phase) was assigned when P was < 2.5 ng/ml, cycle phase 2 (secretory phase) when P was > 5 ng/ml and the LH:FSH ratio was > 1.5 according to the standard of our central laboratory.

### RNA isolation and sequencing

Total RNA from endometrium of rudimentary uterine tissue or normal uterus was isolated using the RNeasy Mini Kit (Qiagen) and used for paired-end RNA-seq. Quality was assessed with an Agilent 2100 Bioanalyzer. Samples with high RNA integrity number (RIN > 7) were selected for library construction. Using the NEBNext Ultra II Directional RNA Library Prep Kit for Illumina and 100 ng of total RNA for each sequencing library, poly(A) selected paired-end sequencing libraries (101 bp read length) were generated according to the manufacturer’s instructions. All libraries were sequenced on an Illumina NovaSeq 6000 platform at a depth of around 40 mio reads each. Library preparation and sequencing procedures were performed by the same individual, and a design aimed to minimize technical batch effects was chosen.

### Quality control, alignment, and differential expression analysis

Read quality of RNA-seq data in fastq files was assessed using *FastQC* (v0.11.4) (38) to identify sequencing cycles with low average quality, adaptor contamination, or repetitive sequences from PCR amplification. Reads were aligned using *STAR* (v2.7.0a) (39) allowing gapped alignments to account for splicing against the *Ensembl* H. sapiens genome v95. Alignment quality was analyzed using *samtools* (v1.1) (40). Normalized read counts for all genes were obtained using *DESeq2* (v1.26.0) (41). Transcripts covered with less than 50 reads (median of all samples) were excluded from the analysis leaving 15,131 genes for determining differential expression. Surrogate variable analysis (sva, v3.34.0) was used to minimize unwanted variation between samples (42). We set |log_2_ fold-change| ≥ 0.5 and BH-adjusted *p*-value ≤ 0.05 to call differentially expressed genes. Gene-level abundances were derived from *DESeq2* as normalized read counts and used for calculating the log_2_-transformed expression changes underlying the expression heatmaps for which ratios were computed against mean expression in control samples. The *sizeFactor*-normalized counts provided by *DESeq2* also went into calculating nRPKMs (normalized Reads Per Kilobase per Million total reads) as a measure of relative gene expression (43). The *sizeFactors* further served in scaling transcript isoform abundances derived from *Salmon* (v0.11.4) (44).

### Gene annotation, enrichments, and regulator analyses

*G:Profiler2* (v0.1.7) was employed to identify overrepresented Gene Ontology terms for differentially expressed genes (45). Upstream regulators as well as predicted interactions among DEGs were derived from *Ingenuity Pathway Analysis* (IPA, v01–16, Qiagen). *Cytoscape* was used for visualizing networks (46). Transcription factor binding site analyses were carried out in *Pscan* (v1.4) (47) on the H. sapiens genome considering -450 to +50 bp of promoter regions for motifs against the JASPAR 2018_NR database. *TFEA.chip* (v1.6) was employed with default parameters to determine transcription factor enrichments using the initial database version of ChIP-Seq experiments (19). Cell type-specific endometrial marker genes were taken from a preprint (13).

### Co-expression analysis

Weighted Gene Co-expression Network Analysis (20) was used to identify gene co-expression. WGCNA is based on the pairwise correlation between all pairs of genes in the analyzed data set. As correlation method, biweight midcorrelation (48) was used with *maxPOutliers* = 0.1, thereby minimizing the influence of potential outliers. Correlations were transformed in a signed hybrid similarity matrix where negative and zero correlations equal zero, while positive correlations remain unchanged. This similarity matrix was raised to the power *β* = 7 to generate the network adjacency and thereby suppressing low correlations that likely reflect noise in the data. For a measure of interconnectedness, adjacency was transformed into a topological overlap measure (TOM) that is informed by the adjacency of every gene pair plus the connection strength they share with neighboring genes. 1-TOM was then given as an input to hierarchical clustering which identified modules, i.e. groups of co-expressed genes by applying the *Dynamic Tree Cut* algorithm (49). Each of these modules was summarized by its first principal component referred to as its eigengene, providing a single value for a module’s expression profile. In order to identify modules affected in MRKH, eigengenes were correlated with the disease trait. A joint Bayesian-frequentistic algorithm combining Bayes Factor (BF) (50) and significance of a correlation was used to identify modules associated with disease status. Modules with an eigengene-trait correlation of *p*_Bonferroni_ = 0.05|BF < 3 were considered significantly associated with MRKH.

## Acknowledgements

We thank all patients who participated in the study.

This study was supported by a project grant of the Deutsche Forschungsgemeinschaft (DFG; BR 5143/5-1, AOBJ: 639534; KO 2313/7-1, AOBJ: 639535; RI 682/15-1, AOBJ: 639536) and through funding of the NGS Competence Center Tübingen (NCCT-DFG, project 407494995). JSH was funded by Brigitte Schlieben-Lange-program from the state of Baden Württemberg.

## Author contribution

SYB, OR, and, OK conceived and designed the project. AK and KR collected and processed patient samples. TH, NW, and JSH analyzed the data and developed the online tool. AKi and SB helped with the analyses. NC was responsible for library preparation and sequencing the samples. TH and JSH wrote the paper. All authors contributed to the interpretation of results and provided critical feedback on preparation of the manuscript.

## Competing interests

The authors declare no competing interests.

## Data availability

RNA-seq data that support the findings of this study have been deposited in the European Genome-phenome Archive (EGA) under primary accession [pending submission validation]. Processed data files are available via the online tool: http://mrkh-data.informatik.uni-tuebingen.de

