## Supplemental Material for "The endometrial transcription landscape of MRKH syndrome"

### Supplementary Figure 1

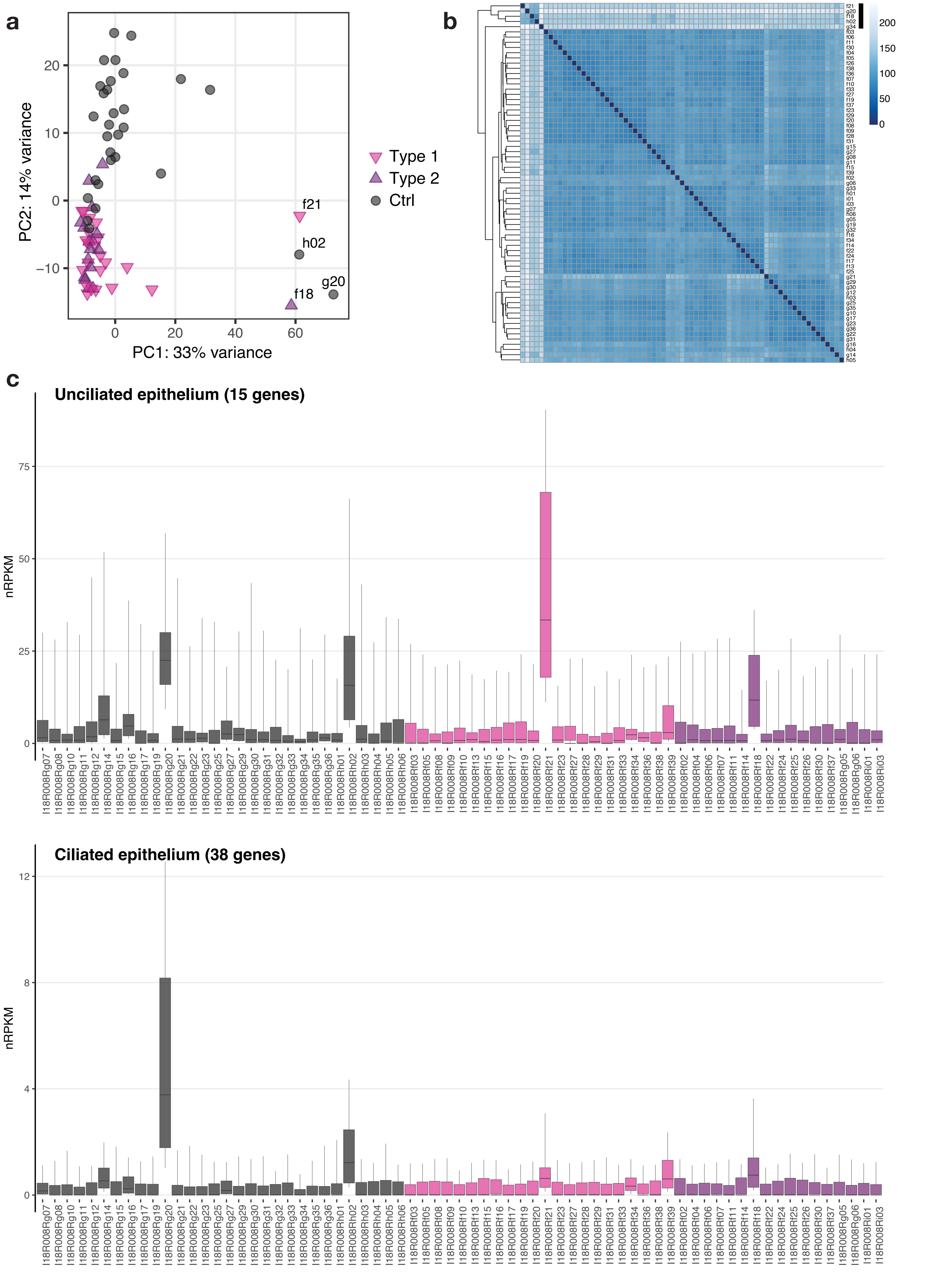

***Supplementary Figure 1: Outlier samples characterized by heterogeneity in cell-type composition.***

a) Principal component analysis of endometrial gene expression profiles for all initial 69 samples. Axis percentages indicate variance contribution of principle components.

b) Correlation map depicting hierarchical clustering of sample-to-sample distances for all initial 69 samples. Variance-stabilized transformed RNA-seq read counts for whole transcriptomes were used to calculate sample-to-sample Euclidean distances (color scale). Results hierarchically clustered using the *complete* method.

c) Cell type-specific gene expression per sample for unciliated and ciliated epithelium. Boxplots show geometric mean as well as 10th, 25th, 75th, and 90th quantile of expression values for all genes classified based on single-cell data of the human endometrium (13). Number of genes per cell type in brackets.

### Supplementary Figure 2

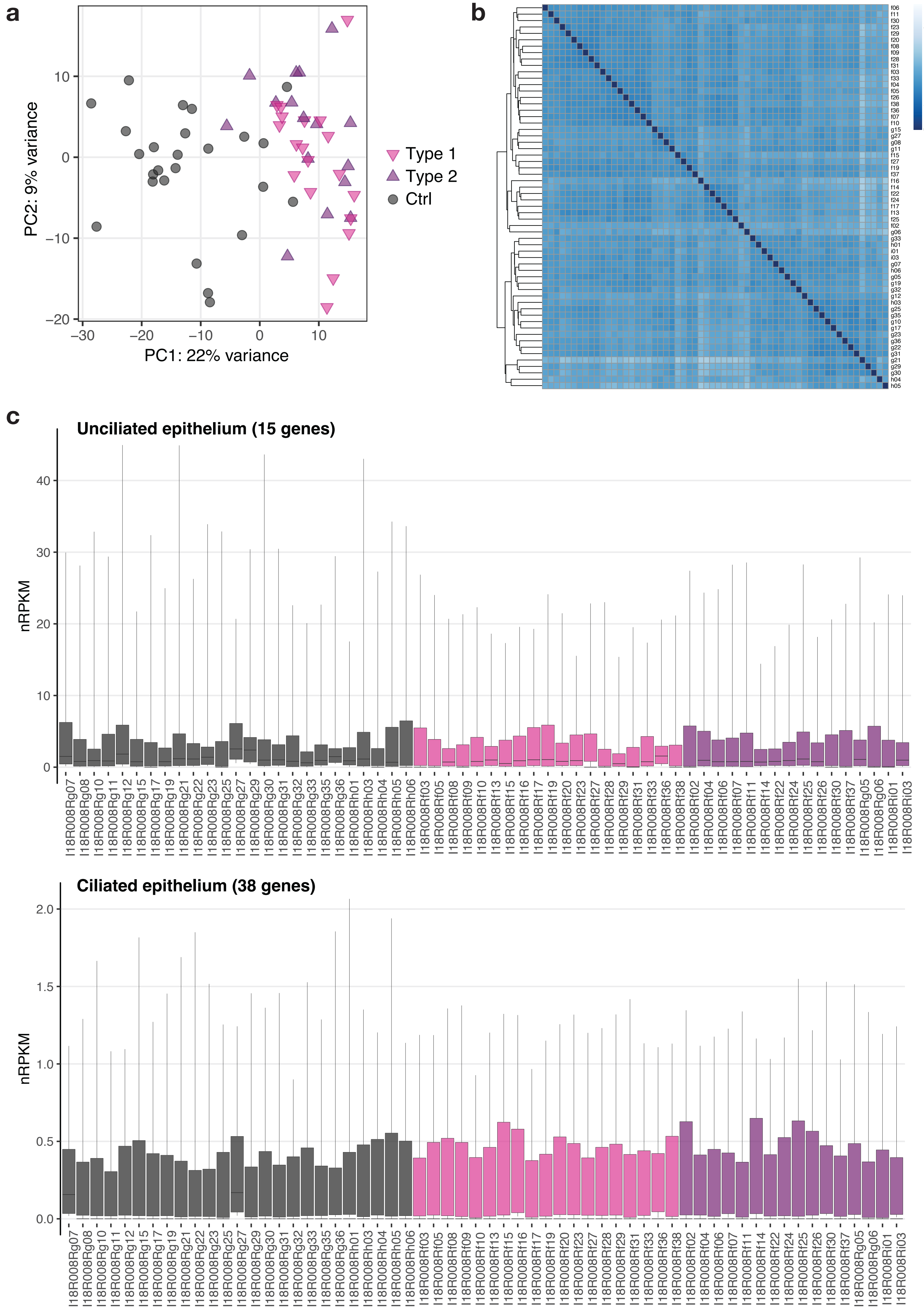

***Supplementary Figure 2: MRKH type 1 and 2 samples separated from control tissue in unaffected individuals.***

a) Principal component analysis of endometrial gene expression profiles for 60 samples after removal of outliers identified in Supplementary Figure 1. Axis percentages indicate variance contribution.

b) Correlation map depicting hierarchical clustering of sample-to-sample distances for the 60 samples remaining after outlier removal. Variance-stabilized transformed RNA-seq read counts for whole transcriptomes were used to calculate sample-to-sample Euclidean distances (color scale). Results hierarchically clustered using the complete method.

c) Cell type-specific gene expression per sample for unciliated and ciliated epithelium. Boxplots show geometric mean as well as 10th, 25th, 75th, and 90th quantile of expression values for all genes classified based on single-cell data of the human endometrium (13). Number of genes per cell type in brackets.

### Supplementary Figure 3

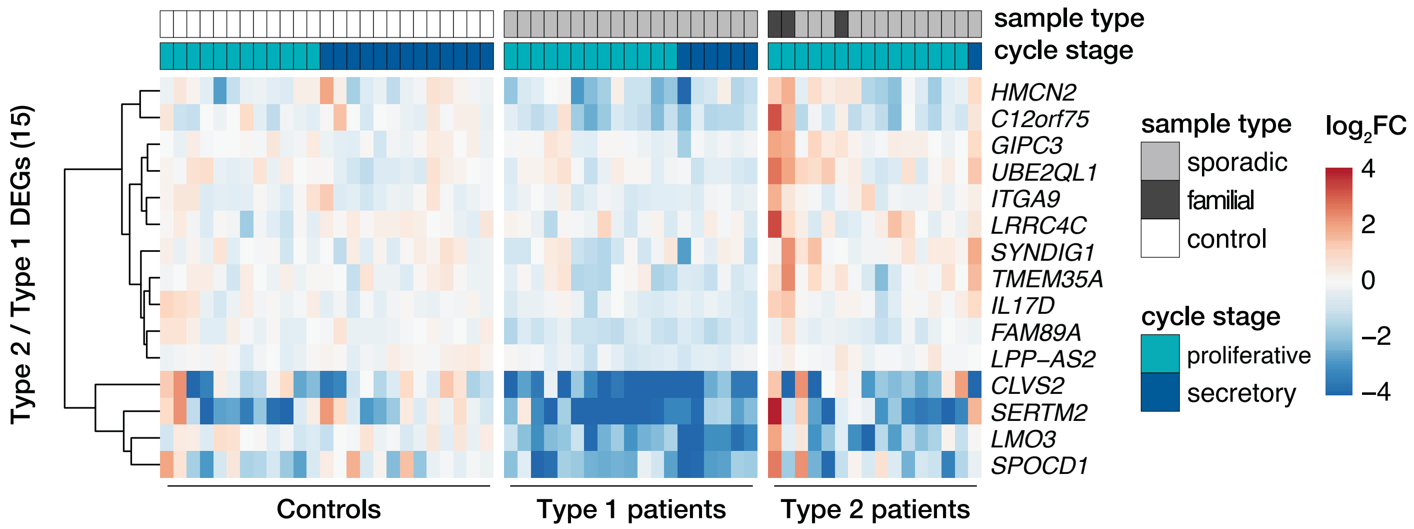

***Supplementary Figure 3: Genes differentially expressed between MRKH type 1 and 2.***

Expression profiles ($log_2$ expression change relative to Ctrl group) of 15 DEGs differentially expressed between MRKH type 1 and type 2 across all samples. Rows hierarchically clustered by Euclidian distance and *ward.D2* method. Cycle information (proliferative or secretory) and patient type (sporadic, familial, or control) on top. For details see Supplementary Table 1.

### Supplementary Figure 4

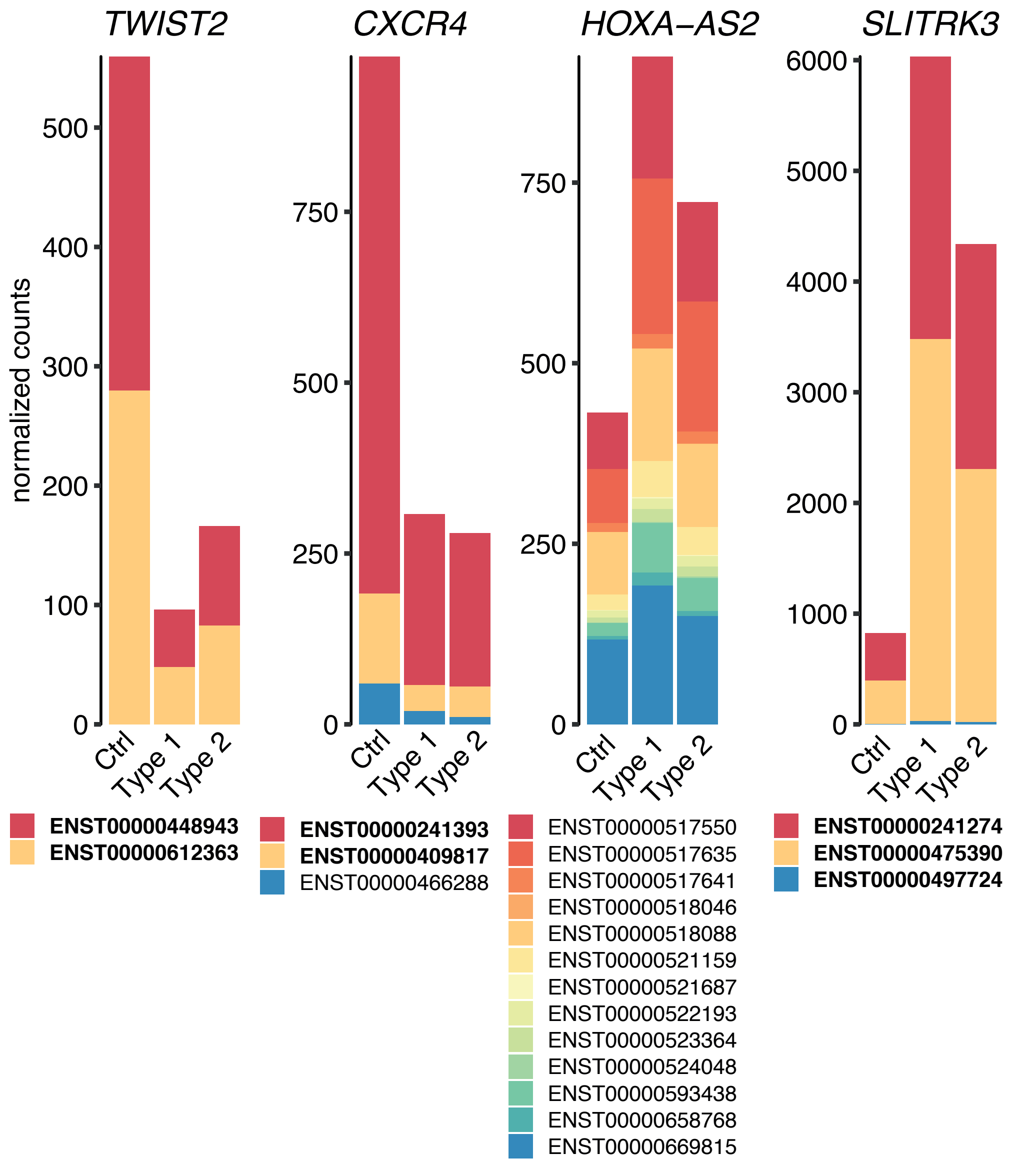

***Supplementary Figure 4: Most significantly altered DEGs showed similar expression changes on transcript isoform level for both types of MRKH.***

Transcript isoform-specific expression changes of *TWIST2*, *CXCR4*, *HOXA-A2*, and *SLITRK3* across all conditions. Mean normalized read counts plotted; bold isoforms are protein-coding.

#### Supplementary Figure 5

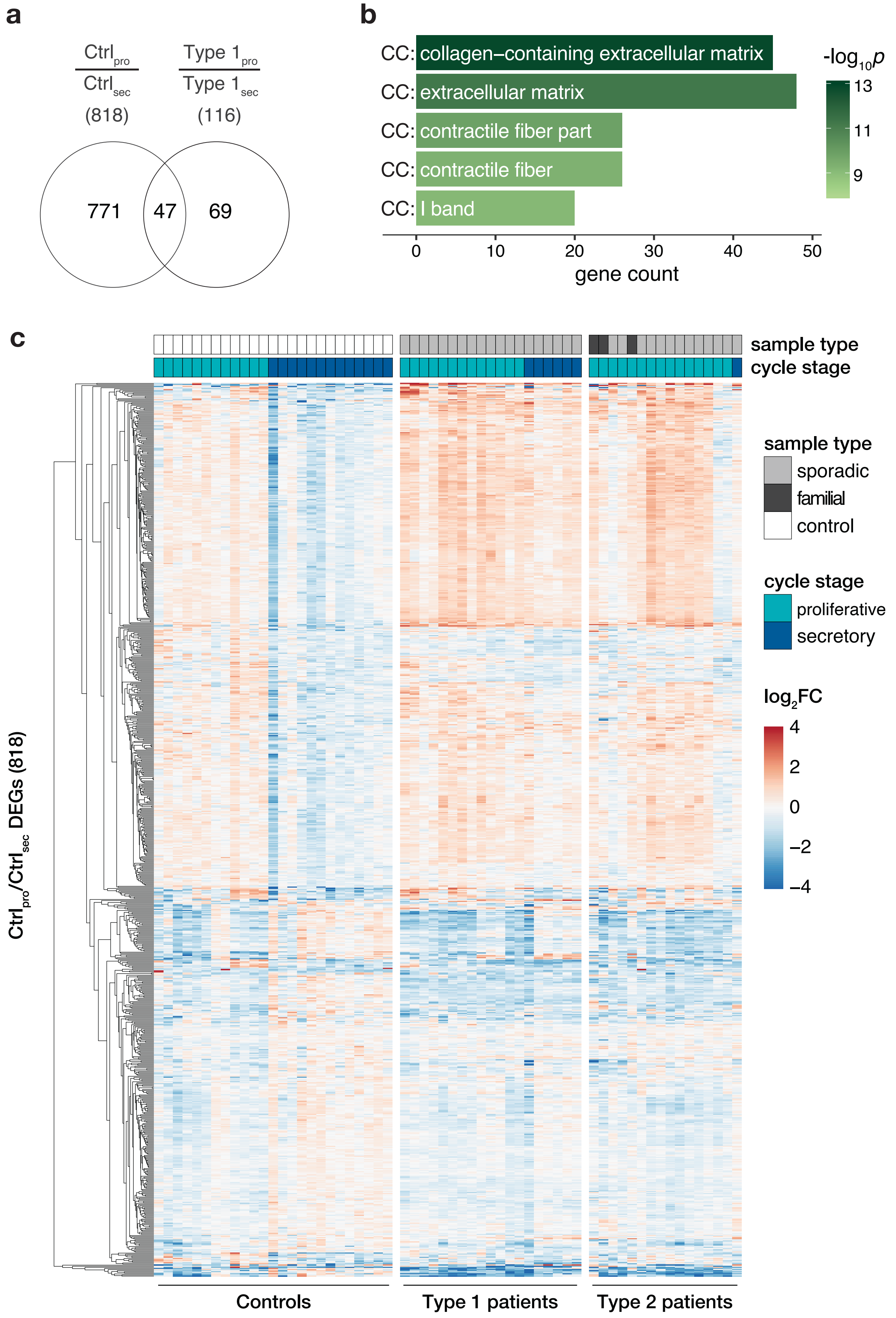

***Supplementary Figure 5: Gene expression changes during the menstrual cycle are missing in endometrium of MRKH patients***

**a)** Venn diagram comparing menstrual cycle-dependent DEGs between control and MRKH type 1 samples.

**b)** Enrichment analysis identified several significantly overrepresented Gene Ontology terms among the 818 DEGs from the control samples (see a). Top five terms with number of associated genes are shown according to their significance. CC: cellular compartment.

**c)** Expression profiles ($log_2$ expression change relative to Ctrl group) of 818 DEGs differentially expressed between proliferative and secretory controls shown across all samples. Rows hierarchically clustered by Euclidian distance and ward.D2 method. Cycle information (proliferative or secretory) and patient type (sporadic, familial, or control) on top. For details see Supplementary Table 1.

### Supplementary Figure 6

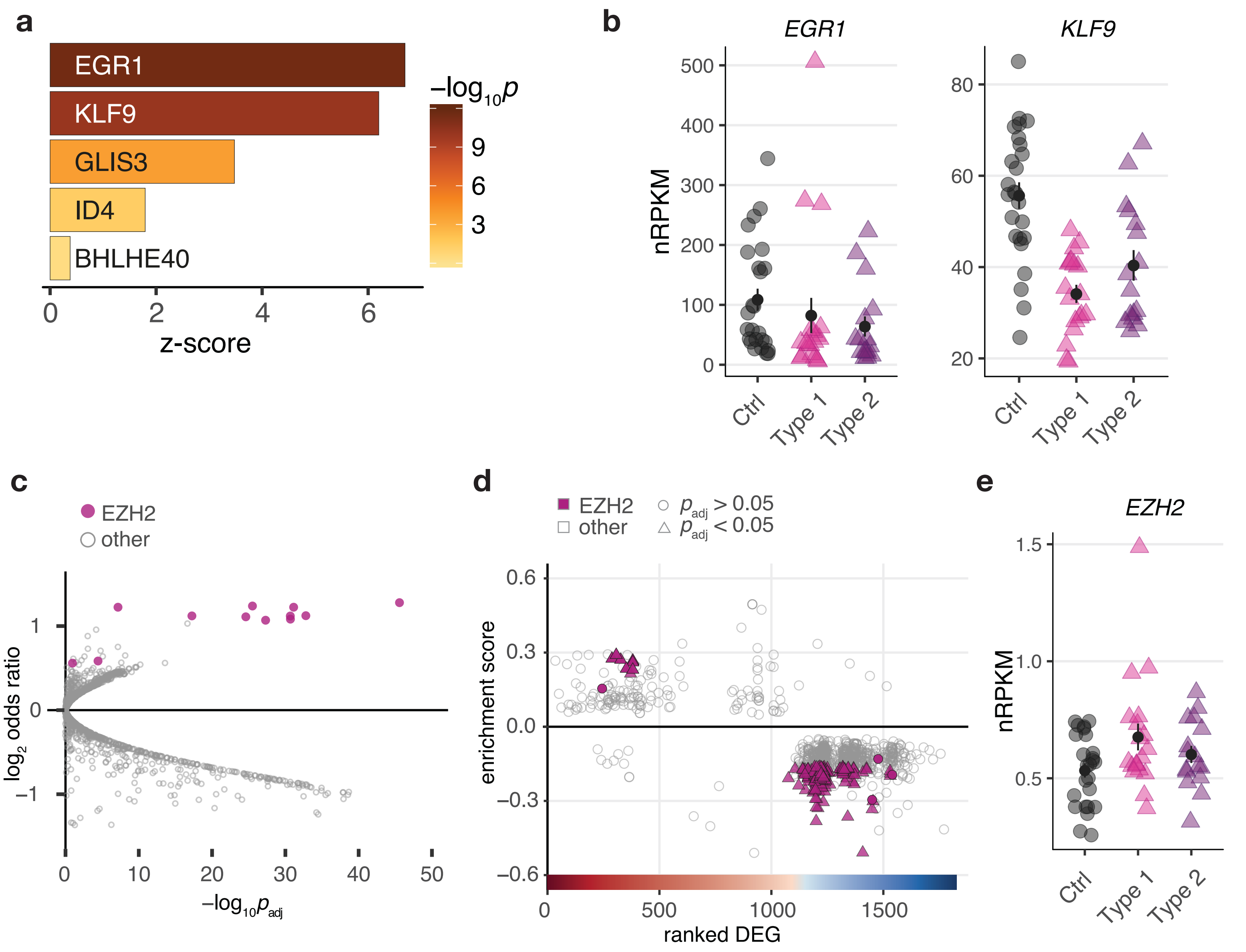

***Supplementary Figure 6: Differentially expressed genes in both types of MRKH are enriched for EGR1, KLF9, and EZH2 binding sites.***

- **a)** Promoter analysis of 2121 DEGSs for overrepresented transcription factor binding sites. Depicted are z-scores for different position weight matrices of known transcription factors that are themselves among the DEG set. Higher z-score reflect higher enrichment. Shown are the top five significantly enriched matrices.
- **b)** Expression levels in normalized reads per kilobase per million (nRPKMs) for *EGR1* and *KLF9* plotted as individual data points with mean ± SEM.
- **c)** Enrichment analysis of transcriptional regulators for the 2121 DEGs identified in MRKH patients based on ChIP-seq and DNase-I data according to TFEA.ChIP. EZH2 is predicted to bind significantly more genes of the DEG set than in the rest of the genome. Analysis based on default parameters for binding sites <1kB upstream including enhancer elements. Each dot represents a ChIP-seq accession, EZH2-related accessions in purple.
- **d)** GSEA-like analysis of transcription regulators for 2121 DEGs based on TFEA.ChIP with default parameters and binding sites <1kB upstream including enhancer elements. Results indicate EZH2 to be the most significantly enriched regulator in the ranked gene set of DEGs.
- **e)** Expression levels in normalized reads per kilobase per million (nRPKMs) for EZH2 plotted as individual data points with mean ± SEM.

#### Supplementary Figure 7

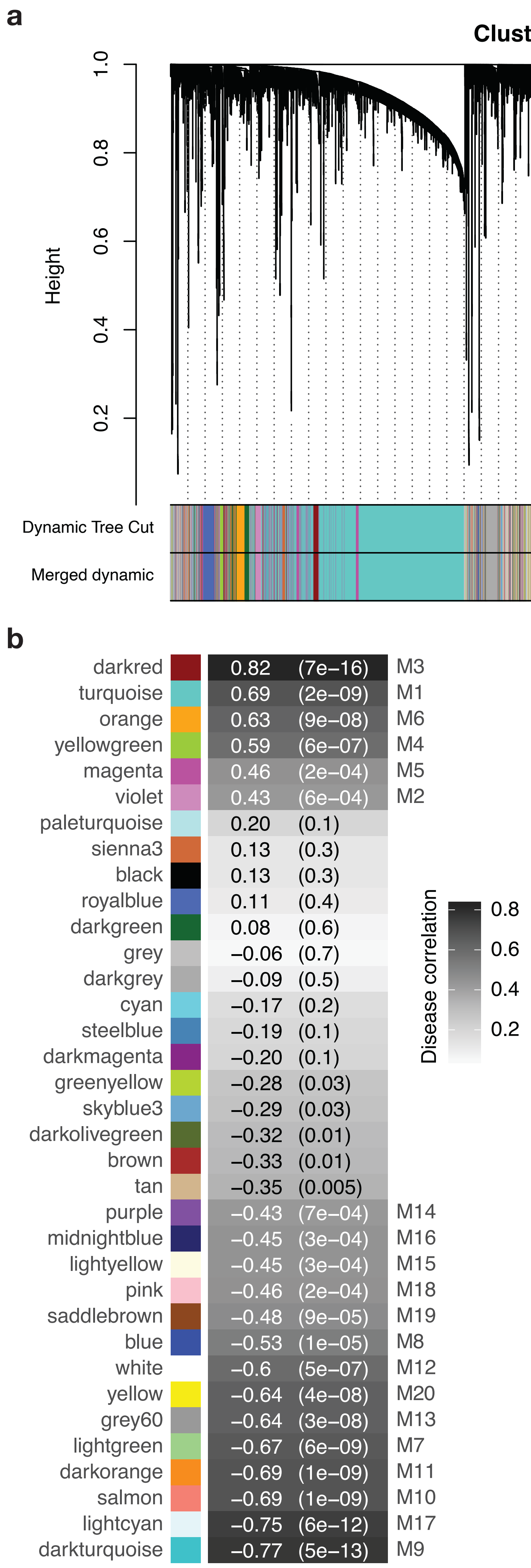

***Supplementary Figure 7: Co-expression analysis points to disease-associated modules.***

- **a)** Representation of the hierarchical gene clustering tree leading to 35 modules of co-expressed genes. Leaves correspond to 15,361 genes included in the analysis, and height reflects closeness of individual genes. Lower panel shows colors assigned to each module by the *Dynamic Tree Cut* and modules assigned consecutive to the *Merged Dynamic* method using a dissimilarity threshold of 0.1.
- **b)** Disease correlations with the corresponding *p*-values for each module. Each cell contains the correlation coefficient between the expression profile of a module and the disease (MRKH) and is color-coded accordingly. Number in brackets indicates associated *p*-value.

**Supplementary Figure 8**

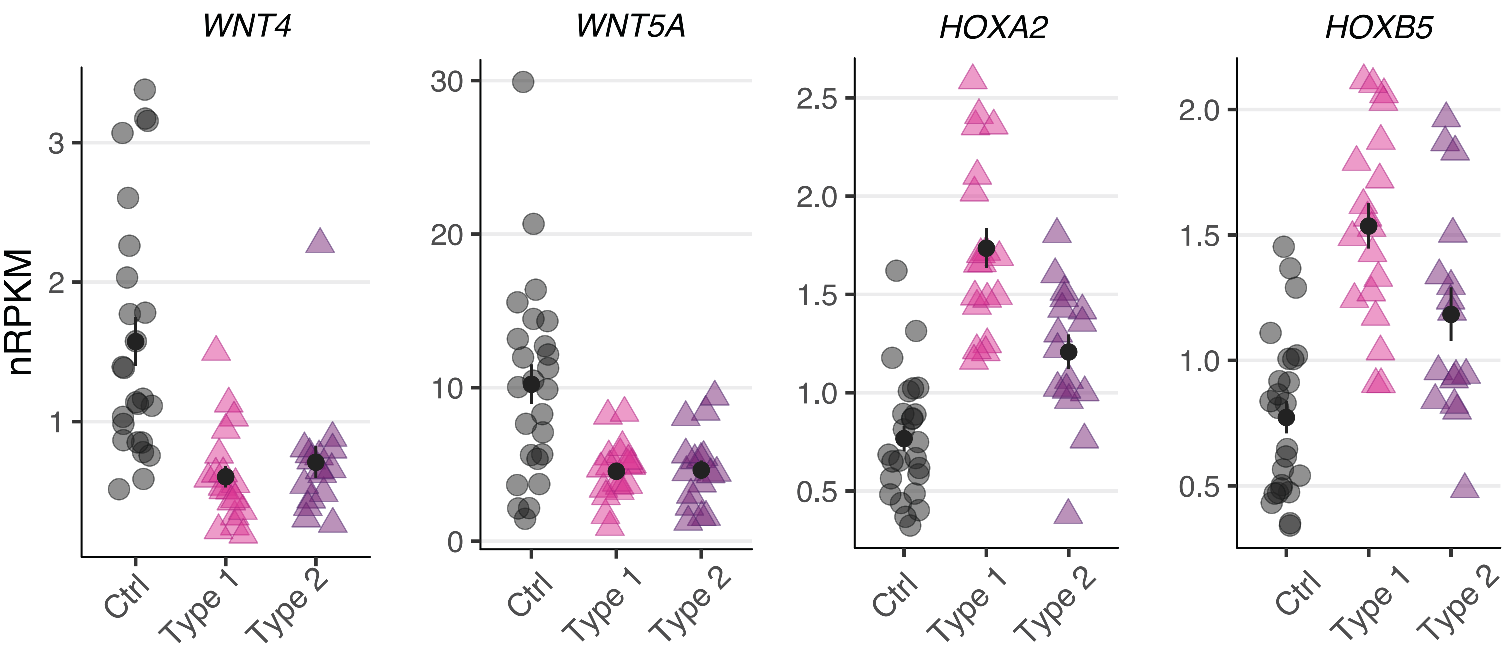

***Supplementary Figure 8: Gene expression changes for altered candidates of the WNT signaling pathway and HOX cluster.***

Expression levels in normalized reads per kilobase per million (nRPKMs) for differentially expressed genes of the WNT signaling pathway and HOX clusters previously linked to MRKH and the development of the Müllerian duct (7). Plotted as individual data points with mean ± SEM.

**Supplementary Table 1.** Demographic and clinical characteristics of MRKH patients and controls

| | Age at surgery (years) | MRKH type | MURCS | Co-occuring anomalies | Cycle phase |
|---|---|---|---|---|---|
| I18R008Rg07 | 37 | N/A | N/A | N/A | 2 |
| I18R008Rg08 | 33 | N/A | N/A | N/A | 1 |
| I18R008Rg10 | 35 | N/A | N/A | N/A | 2 |
| I18R008Rg11 | 35 | N/A | N/A | N/A | 1 |
| I18R008Rg12 | 35 | N/A | N/A | N/A | 1 |
| I18R008Rg14 [Δ] | 34 | N/A | N/A | N/A | 2 |
| I18R008Rg15 | 36 | N/A | N/A | N/A | 1 |
| I18R008Rg16 [Δ] | 35 | N/A | N/A | N/A | 2 |
| I18R008Rg17 | 35 | N/A | N/A | N/A | 2 |
| I18R008Rg19 | 34 | N/A | N/A | N/A | 1 |
| I18R008Rg20 [Δ] | 34 | N/A | N/A | N/A | 2 |
| I18R008Rg21 | 37 | N/A | N/A | N/A | 2 |
| I18R008Rg22 | 37 | N/A | N/A | N/A | 2 |
| I18R008Rg23 | 35 | N/A | N/A | N/A | 2 |
| I18R008Rg25 | 31 | N/A | N/A | N/A | 1 |
| I18R008Rg27 | 34 | N/A | N/A | N/A | 1 |
| I18R008Rg29 | 35 | N/A | N/A | N/A | 2 |
| I18R008Rg30 | 37 | N/A | N/A | N/A | 2 |
| I18R008Rg31 | 37 | N/A | N/A | N/A | 2 |
| I18R008Rg32 | 37 | N/A | N/A | N/A | 1 |
| I18R008Rg33 | 36 | N/A | N/A | N/A | 1 |
| I18R008Rg34 [Δ] | 34 | N/A | N/A | N/A | 2 |
| I18R008Rg35 | 34 | N/A | N/A | N/A | 1 |
| I18R008Rg36 | 36 | N/A | N/A | N/A | 2 |
| I18R008Rh01 | 37 | N/A | N/A | N/A | 2 |
| I18R008Rh02 [Δ] | 37 | N/A | N/A | N/A | 1 |
| I18R008Rh03 | 38 | N/A | N/A | N/A | 2 |
| I18R008Rh04 | 38 | N/A | N/A | N/A | 1 |
| I18R008Rh05 | 38 | N/A | N/A | N/A | 2 |
| I18R008Rh06 | 38 | N/A | N/A | N/A | 1 |
| I18R008Rf03 | 18 | 1 | No | None | 1 |
| I18R008Rf05 | 16 | 1 | No | None | 1 |
| I18R008Rf08 | 16 | 1 | No | None | 2 |
| I18R008Rf09 | 19 | 1 | No | None | 2 |
| I18R008Rf10 | 17 | 1 | No | None | 1 |
| I18R008Rf13 | 18 | 1 | No | None | 1 |
| I18R008Rf15 | 17 | 1 | No | None | 1 |
| I18R008Rf16 | 17 | 1 | No | None | 2 |
| I18R008Rf17 | 16 | 1 | No | None | 1 |
| I18R008Rf19 | 24 | 1 | No | None | 1 |
| I18R008Rf20 | 15 | 1 | No | None | 2 |
| I18R008Rf21 [Δ] | 18 | 1 | No | None | 1 |
| I18R008Rf23 | 24 | 1 | No | None | 1 |
| I18R008Rf27 | 19 | 1 | No | None | 1 |
| I18R008Rf28 | 19 | 1 | No | None | 2 |
| I18R008Rf29 | 16 | 1 | No | None | 1 |
| I18R008Rf31 | 17 | 1 | No | None | 2 |

| | | | | | |
|---|---|---|---|---|---|
| I18R008Rf33 | 15 | 1 | No | None | 1 |
| I18R008Rf34 [Δ] | 16 | 1 | No | None | Unknown |
| I18R008Rf36 | 17 | 1 | No | None | 1 |
| I18R008Rf38 | 14 | 1 | No | None | 1 |
| I18R008Rf39 [Δ] | 17 | 1 | No | None | 1 |
| I18R008Rf02* | 30 | 2 | Yes | Duplex kidney (R), scoliosis | 1 |
| I18R008Rg05* | 45 | 2 | Yes | Duplex kidney (B), complex sceletal malformations | 1 |
| I18R008Rf18° [Δ] | 16 | 2 | No | Scoliosis | 1 |
| I18R008Rg06° | 16 | 2 | No | Scoliosis | 1 |
| I18R008Rf04 | 16 | 2 | No | Duplex kidney (B) | 1 |
| I18R008Rf06 | 20 | 2 | No | Scoliosis | 1 |
| I18R008Rf07 | 21 | 2 | No | Ear deformities, auditory canal hypoplasia | 1 |
| I18R008Rf11 | 16 | 2 | No | Scoliosis | 1 |
| I18R008Rf14 | 19 | 2 | No | Renal agenesis (R) | 1 |
| I18R008Rf22 | 17 | 2 | No | Scoliosis | 1 |
| I18R008Rf24 | 23 | 2 | No | Scoliosis | 1 |
| I18R008Rf25 | 18 | 2 | Yes | Renal agenesis (L), scoliosis, ear deformities, fistula, imperforate anus | 1 |
| I18R008Rf26 | 17 | 2 | No | Renal agenesis (R) | 1 |
| I18R008Rf30 | 21 | 2 | No | Hip dysplasia | 2 |
| I18R008Rf37 | 25 | 2 | No | Silver-Russell syndrome | 1 |
| I18R008Ri01 | 18 | 2 | No | Renal agenesis (R) | 1 |
| I18R008Ri03 | 18 | 2 | No | Nephroptosis (L) | 1 |

[Δ] patient samples excluded from analysis due to deviating tissue composition; * two sisters from family A; ° two sisters from family B; N/A , not applicable (hysterectomy control); B = bilateral, L = left, R= right; cycle phase 1 ≙ proliferative phase, cycle phase 2 ≙ secretory phase

**Supplementary Table 2.** Summary of demographic and clinical characteristics of MRKH patients and controls used for the analysis. Samples excluded from the analysis due to tissue composition (see Supplementary Table 1) were not included in the calculation of the depicted values.

| | No of patients | Age at surgery (years), mean ± Stdev | % MURCS | % Renal malformations | % Sceletal malformations | % Further malformations | Cycle phase | |
|---|---|---|---|---|---|---|---|---|
| | | | | | | | % phase 1 | % phase 2 |
| Controls | 25 | 35.8 ± 1.72 | N/A | N/A | N/A | N/A | 48,00 | 52,00 |
| MRKH | 35 | 19.26 ± 5.5 | 8,57 | 22,86 | 28,57 | 8,57 | 80,00 | 20,00 |
| MRKH type 1 | 19 | 17.58 ± 2.58 | 0,00 | 0,00 | 0,00 | 0,00 | 68,42 | 31,58 |
| MRKH type 2 | 16 | 21.25 ± 7.14 | 18,75 | 50,00 | 62,50 | 18,75 | 93,75 | 6,25 |

N/A, not applicable (hysterectomy control); cycle phase 1 ≙ proliferative phase, cycle phase 2 ≙ secretory phase
